# Denoising 7T Structural MRI with Conditional Generative Diffusion Models

**DOI:** 10.1101/2025.07.03.663033

**Authors:** Yixin Wang, Binxu Li, Yihao Liang, Mackenzie Carlson, Phillip DiGiacomo, Hossein Moein Taghavi, Julian Maclaren, Meghan Bell, Nancy Pham, Johannes Hugo Decker, Michael lv, Tie Liang, Jason Wong, Elizabeth Mormino, Victor W. Henderson, Greg Zaharchuk, Brian Rutt, Wei Shao, Marios Georgiadis, Michael Zeineh

## Abstract

**Purpose:** 7T MRI offers ultra-high resolution and improved sensitivity for iron deposition in neurodegenerative disorders, but commonly used acquisitions are long and hence challenging, especially for elderly subjects. Efficiently denoising a short acquisition to achieve the image quality of a longer acquisition would be of translational benefit.

**Materials and Methods:** We introduce a conditional diffusion model derived from generative AI (a 7T Conditional Diffusion Model, 7TCDM) that was trained on native single-acquisition 2D reconstructions and referenced multi-repetition images to guide the denoising process and improve SNR and contrast. 7TCDM model was tested on 2D T2-weighted gradient-echo imaging from 19 participants, including healthy controls and individuals with mild cognitive impairment or Alzheimer’s disease (AD). 7TCDM’s performance was assessed using Mean Squared Error (MSE), Peak Signal-to-Noise Ratio (PSNR), Structural Similarity Index Measure (SSIM), and comprehensive reader studies.

**Results:** Referencing the multi-repetition ground truth, 7TCDM improved the single-acquisition original image by 29.1% in MSE, 5.8% in PSNR, and 9.4% in SSIM, and outperformed convolutional neural network-based models in all metrics. Expert rater evaluations confirmed superior image quality, with significantly enhanced detail and contrast preservation in regions such as the hippocampi, white matter lesions, and small cortical veins. The model also demonstrated robust performance in both the concurrently acquired and publicly available 3D multi-echo gradient echo acquisitions, which the model was not trained on.

**Conclusions:** The 7T Conditional Diffusion Model provides high-quality denoised images from shorter scans, increasing the feasibility of scanning patients in shorter times while preserving essential anatomical and pathological details.

## 1 Introduction

Magnetic resonance imaging (MRI) is a crucial noninvasive diagnostic tool in modern healthcare. Ultrahigh-field MRI at 7 Tesla (7T MRI) provides significant advantages over lower magnetic fields due to its higher signal-to-noise ratio (SNR) and contrast-to-noise ratio (CNR) for magnetic susceptibility ^1^. The enhanced SNR/CNR can be translated to increased resolution and lesion contrast, which dramatically improves the conspicuity of structural changes within the brain, including microscopic iron deposition that has previously only been detectable through post-mortem tissue analyses, such as in Alzheimer’s disease ^2,3^.

However, 7T MRI introduces challenges such as B0 and B1 inhomogeneity, susceptibility-related artifacts, and chemical shift errors, which degrade image quality by introducing noise and artifacts ^4^. Furthermore, to take advantage of improved SNR/CNR, 7T sequences are typically very high-resolution and often long in duration, resulting in increased motion artifact in vulnerable groups such as the elderly ^5,6^. In research settings, one approach to increase SNR and minimize the contribution of artifacts is to perform multiple sequential repetitions and average them to obtain a higher quality image to depict anatomic details and pathology ^7^. However, this averaging has significant downsides: each repetition takes several minutes, cumulatively leading to long scan times. Also, with very high resolution, simple signal averaging is not possible due to small amounts of patient motion. Therefore, denoising a scan with fewer repetitions could more efficiently solve the averaging challenges by minimizing artifacts and noise.

Deep learning models have shown remarkable success in medical imaging denoising. Previous studies applied Convolutional Neural Network (CNN)-based models ^8^ to MRI ^9,10^, now adopted clinically/commercially. However, CNN-based models tend to directly learn a mapping between the input and output images, with fine details often lost as a result of over-smoothing ^11^. An alternative strategy is Generative Adversarial Networks (GANs) for image denoising ^12^. However, GANs are prone to unstable training, especially with the limited sample sizes typical for 7T MRI ^13^. Current approaches to 7T MRI reconstruction and denoising have typically relied on traditional optimization ^14,15^ or have focused on small-scale applications like mouse brain imaging using CNN-based models ^16^. Our work aims to address the limitations of current techniques by obtaining high-quality 7T MRI images from a single repetition without sacrificing crucial details using advanced generative artificial intelligence (AI) techniques.

In this study, we built a reconstruction model (7T Conditional Diffusion Model [7TCDM]) based on the cutting-edge generative AI model, Denoising Diffusion Probabilistic Model (DDPM) ^17^. DDPMs are a powerful class of generative models that have demonstrated success in multiple medical imaging applications, including low-dose CT denoising ^18^, PET image denoising ^19^, and 1.5T/3T MRI super-resolution tasks ^20,21^. DDPMs learn complex data distributions by using a variational inference framework to effectively estimate the parameters using two core components: a forward diffusion process, which gradually adds noise to the data to map it to a Gaussian distribution, and a reverse diffusion process, which progressively removes the noise to reconstruct the high-quality image. Moreover, we employ “conditional generation” ^17^ to enhance the reconstruction process and preserve structural details. This allows the model to generate images that retain intricate structural information while eliminating noise, resulting in high-quality reconstructions.

Training and testing data consisted of four distinct repetitions of a prospectively motion-corrected 2D gradient-recalled-echo (GRE) sequence ^7^. The model was trained to use one repetition to produce an image matching the average of the four repetitions. It was then tested on participants from our institution’s Alzheimer’s Disease Research Center, which we assessed using quantitative metrics and a blinded neuroradiology reader study. The model also generalized to 3D T2* multi-echo GRE, including publicly available datasets from other magnet vendors ^22^. Codes and models will all be made publicly available.

## 2. Materials and Methods

### 2.1 Image Acquisition

Five healthy volunteers underwent eleven 7T scans (4/3/2/1/1 scans per subject) consisting of 4 repetitions of a 2D gradient-echo sequence with written informed consent and approval of our Institutional Review Board, in accordance with the Health Insurance Portability and Accountability Act of 1996 (Data were also utilized from a previous study ^7^). Images were acquired on a 7T scanner (MR950, GE Healthcare) utilizing prospective motion correction with a 2/32-channel Nova T/R coil. GRE parameters were: voxel size 0.176×0.176×1.0mm^3^, TR=300ms, TE=20ms, FA=35°, six slices spaced every 6mm, 5 min each repetition, four repetitions acquired ^23^. Natively reconstructed images from each repetition were coregistered to account for residual motion using 2D FLIRT ^24^ with sinc interpolation and averaged to obtain a high-quality mean image (hereafter named 4REPS)^23^, which would serve as ground truth for denoising.

Data using the same acquisition were also acquired on 19 participants (5 male and 14 female, age range: 41.5-85.7 years, mean age: 72.6 years) from our institution’s Alzheimer’s Disease Research Center (ADRC), comprising 8 healthy elderly control, 6 subjects with consensus diagnoses of mild cognitive impairment, and 5 with suspected Alzheimer’s disease.

3D multi-echo GRE data for each ADRC subject were also acquired: voxel size 0.43×0.43×1.2mm^3^, TR = 34.78ms, five echo times (TE) at 3.66/10.03/16.40/22.77/29.14ms.

### 2.2 Augmentation and Histogram Matching

To increase the training sample size, each repetition from each training scan was separately coregistered to the other three repetitions (which were in slightly different positions), and the coregistered four repetitions were averaged to create the 4REPS in four different spaces (i.e., the spaces of the 1^st^, 2^nd^, 3^rd^, and 4^th^ repetitions).

Each repetition differed slightly in signal levels due to typical variations at 7T, but all were paired with the same 4REPS, denoted as *x_4REPS_*. To ensure that the model focused on learning the structural reconstruction for each repetition rather than histogram mapping, before training we performed histogram matching between *x_4REPS_* and each single repetition *c*, resulting in a ground truth *x*_0_ (*x*_0_= histogram matched *x_4REPS_*), histogram-matched to a single repetition *c*. Specifically, we matched the cumulative distribution function (CDF) of the histograms of the *H*(*x_4REPS_*) to the CDF of one repetition *G*(*c*) by:

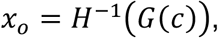

where *H*^−1^ was achieved by one-dimensional interpolation. *x*_0_ will serve as the ground truth and be paired with each repetition *c* to train the models. Each input *c* and *x*_0_ were flipped left-right with a probability of 50% for further augmentation.

### 2.3 Conditional Diffusion Model

7T CDM is built on a generative denoising diffusion model ^19^ for transforming a single repetition image into a 4REPS image, utilizing the single repetition ***c*** as a condition to guide the generation process **(Fig.1).** The diffusion process consists of a forward and reverse process, and both processes are carried out in a multi-step manner, to gradually add and remove noise to synthesize high-quality images.

**Figure 1.**
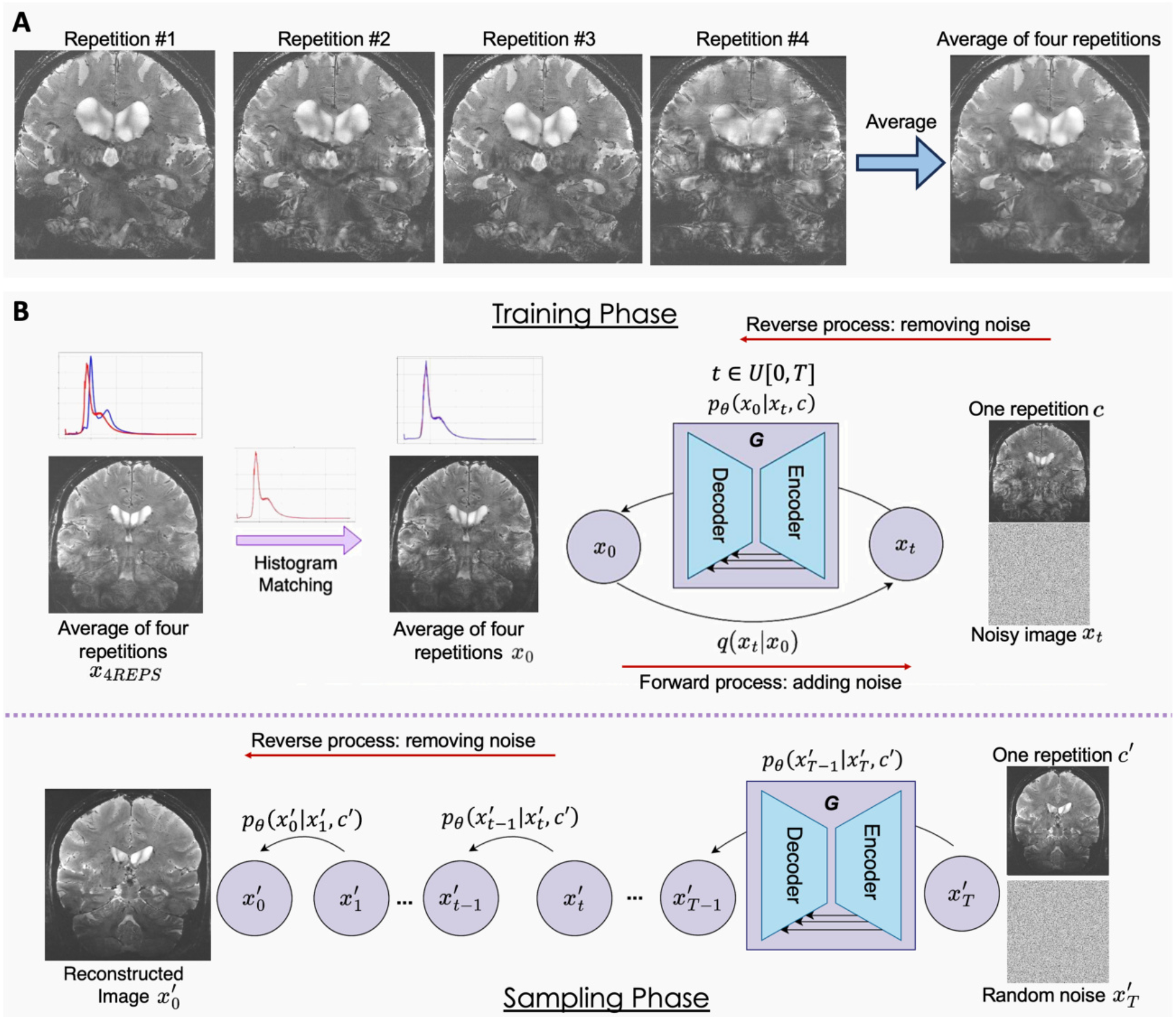
**A.** Illustration of the coregistered four repetitions and their averaged 4REPS, which, after histogram matching, served as the ground truth “denoised” image. **B.** Pipeline of the 7T MR conditional diffusion model (7TCDM). During the training phase, 4REPS was histogram-matched to one single repetition **c** over the imaging volume and converted into **x_0_**. In the forward process, we sampled a timestep **t** from a uniform distribution **U**[**0, T**] and added noise to **x_0_**, resulting in **x_t_**. In the reverse process, the model was trained to approximate the posterior distribution **q**(**x**′**_0_**|**x**′**_t_, c**), reconstructing **x**′**_0_** by removing noise from **x_t_**, conditioned on **c**. After model training was complete and during the sampling phase, the model iteratively removed noise to generate a high-quality image from repetition **c**, and pure Gaussian noise.

#### 2.3.1 Training Phase

Suppose *x*_0_ follows a distribution *q*(*x*_0_):*x*_0_∼*q*(*x*_0_), and *x*_1_,*x*_2_,*x*_3_, …*x*_T_ are noisy data with the same dimension of *x*_0_, where *T* represents the maximum number of time steps, which we set as 10 (See Supplementary Ablation Study). The forward diffusion gradually generates noisy data by adding gaussian noise to the “denoised” data *x*_0_ given timestep *t* ∈*U*[0,*T*], as follows:

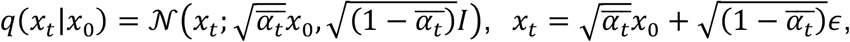

where ε ∼ 𝒩(0,*I*) and 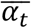 follows cosine-beta schedule^25^ to ensure some noise is added in the beginning.

In the reverse process, we utilized *c* as a condition by concatenating it as an additional channel together with *x_t_*, and trained a model ***G*** with parameters θ to estimate a distribution *p*_0_(*x*_*t*−1_|*x_t_, c*), which approximated the true posterior distribution *q*(*x*_0_|*x_t_, c*). Following Bayes theorem^17^, the reverse process can be formalized as the products of the stepwise posterior probabilities:

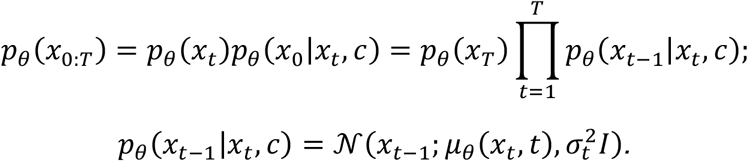

The training objective was to minimize the differences of distribution in forward and reverse processes at the same time step using Kullback-Leibler (KL) divergence. The loss function *L* was written as:

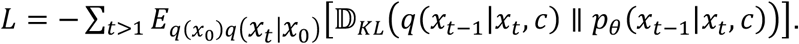

The posterior distribution *q*(*x*_*t*−1_|*x_t_, c*) is Gaussian. Thus, the above objective became the distance between the sampled *x*_*t*−1_ from the posterior and the predicted images, conditioning on time step *t* and *c*, which could be simplified as a Mean Squared Error (MSE) loss function:

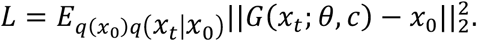

We further forced the model to incorporate image contrast using a Structural-Similarity-Index-Measure (SSIM)^26^ into the objective function, defined as:

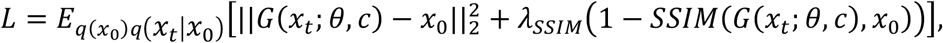

where *λ*_SSIM_ controlled the model’s emphasis on the contrast of the generated images.

#### 2.3.2 Sampling Phase

During the sampling process, the model removed noise from one repetition ***c***^r^ step by step to generate a high-quality image *x*′_0_. We used *p*_0_(*x*′_*t*−1_|*x*′*_t_, c*′) to approximate *q*(*x*′_*t*−1_|*x*′*_t_, c*′) and sampled every step from the previous step, following DDPM ^17^. We used a Denoising Diffusion Implicit Model (DDIM) ^27^ for efficient sampling, where only fewer steps (<= *T*) were required to obtain the final denoised images (see Supplementary Ablation study for inference time). No histogram matching was done in order to match realistic usage.

### 2.4 Model Comparisons

#### Quantitative Comparison

We denoised each single repetition (Original) using 7TCDM (Original: 7TCDM) and compared the performance with Denoising CNN (DnCNN)^8^ (Original: DnCNN), a widely used denoising CNN architecture. Both models were trained on 11 scans from MRIs of five healthy adults (See Supplementary Experimental Settings and Ablation Study).

Inferences were performed on single-repetition images on the 19 held-out ADRC subjects. To see if denoising could be improved by combining with image averaging, the denoised reconstructions of each repetition for each subject using 7TCDM and DnCNN models were averaged to produce a single imaging volume, 4REPS: 7TCDM and 4REPS: DnCNN, respectively.

We compared the reconstruction performance of the 7TCDM and DnCNN models with 4REPS as ground truth using three commonly used metrics: Mean Squared Error (MSE), SSIM, and Peak Signal-to-Noise Ratio (PSNR), all compared with a Wilcoxon rank-sum test, considering each 2D slice as a separate data point, N=19 subjects * 6 slices = 114. To visualize feature loss associated with denoising, we depicted the residuals as the absolute difference between denoised and original images.

Finally, we applied 7TCDM to 2D images extracted from a high-resolution 3D gradient-echo (GRE) acquisition. The 3D data were first unsampled to match the in-plane resolution of the 2D GRE training data. Denoising was then applied directly to each slice using the same model and parameters, with no additional training or tuning required.

#### Rater Analysis

The Original and denoised images were rated at the subject level in a blinded reader study conducted by three experienced neuroradiologists with 3, 7, 14 years of neuroradiology experience using the Horos medical image visualization software. The raters were provided with sample images from the training data and participated in two sessions to facilitate agreement and fully develop a 5-point image quality rating scale (**Supplementary Table 1**): 1 = excellent, 2 = very good, 3 = good, 4 = adequate, and 5 = limited. Following this, the readers then independently and blind to image category compared Original, Original:DnCNN, Original:7TCDM, 4REPS, 4REPS:7TCDM, and 4REPS:DnCNN. Readers were then asked to rank the above six contrasts from 1 to 6, where 1 indicates the best and 6 the lowest. Rater agreements were statistically evaluated by Gwet’s AC^28^ (**Supplementary Fig. 4**). Agreement was high for “RANK ORDER” but not the other assessment components because of mild baseline shifts along the ordinal scale, so categories were averaged across raters, and these means were compared using a Wilcoxon rank-sum test.

## 3. Experimental Results

### 3.1 Quantitative Comparison of Denoising Techniques

Visual improvements in image quality were observed for all DnCNN and especially 7TCDM reconstructions, both on the Original (single repetition) as well as the 4REPS datasets **(Fig.2). Figure 3** shows that Original:7TCDM achieves an MSE of 0.0421±0.0001, an SSIM of 0.876±0.022, and a PSNR of 34.2±1.4 which are significantly higher than both the original images (MSE: 0.0595±0.0002, *p=0.0073*; SSIM: 0.801±0.027, *p<0.0001*; PSNR: 32.6±1.5, *p=0.0089*) and the Original:DnCNN (MSE: 0.0509±0.0002, *p=0.0073*; SSIM: 0.861±0.027, *p<0.0001*; PSNR: 33.3±1.5, *p=0.0089*). The residuals between the original images and 4REPS and denoised images further show DnCNN discards potentially useful structural information, while the 7TCDM residual images contain relatively less structural detail (Supplementary Residual Comparisons).

**Figure 2.**
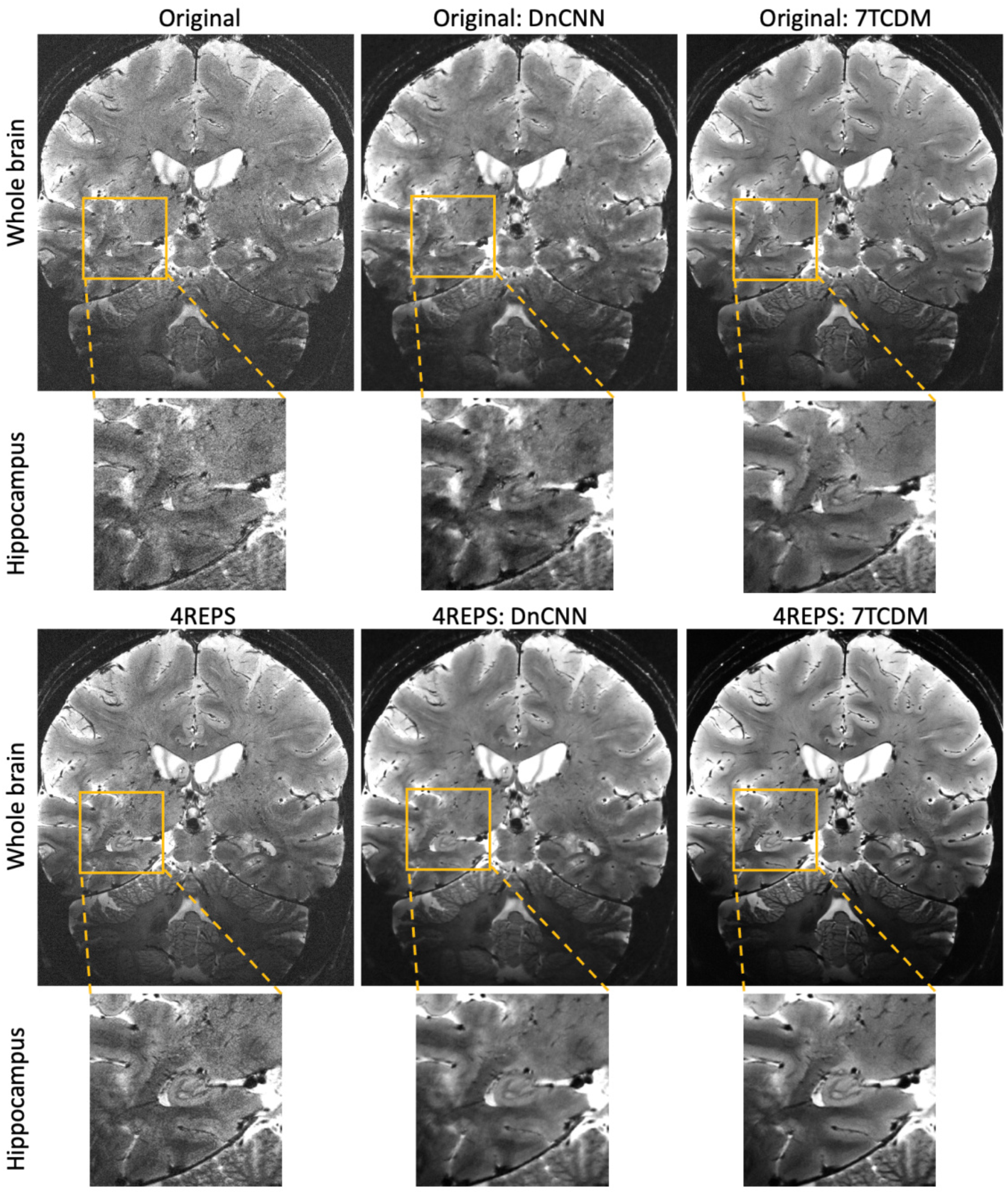
Visual comparisons between the original image from a single repetition and its corresponding denoised images generated by DnCNN (Original:DnCNN) and 7TCDM (Original:7TCDM) on the whole brain (first row) and zoomed-in hippocampus region (second row). The third and fourth rows show the average of four repetitions (4REPS) and the average of denoised images from four repetitions produced by DnCNN (4REPS:DnCNN) and 7TCDM (4REPS:7TCDM), respectively.

**Figure 3.**
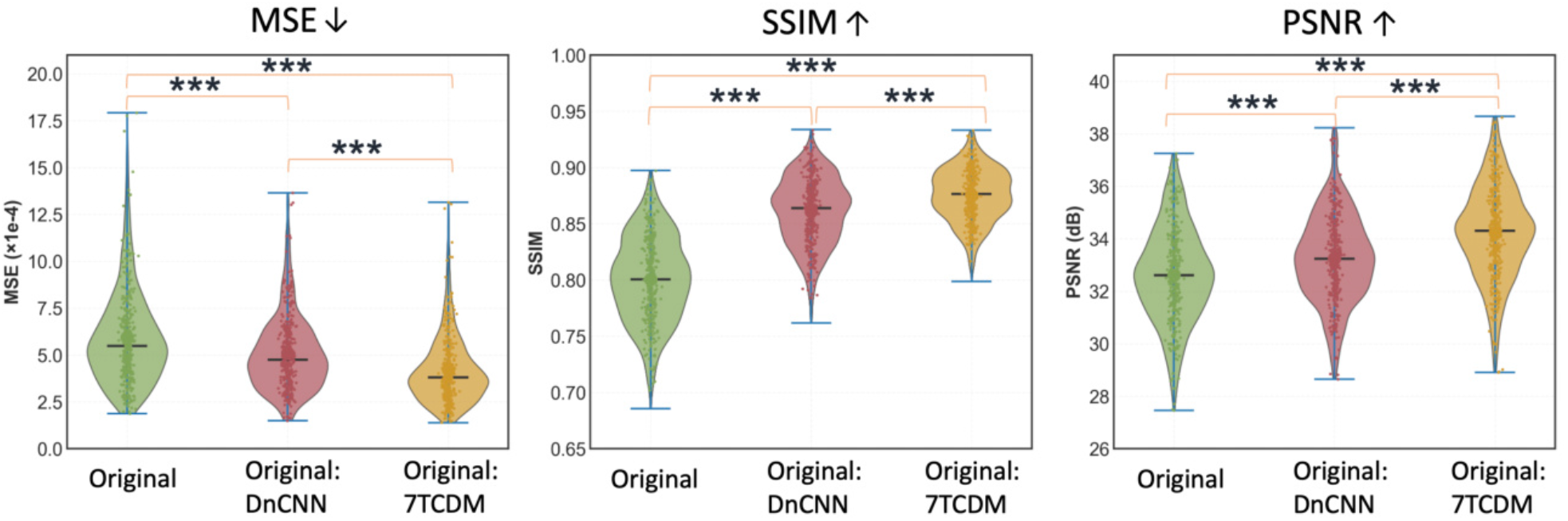
Violin plots of Mean-Squared-Error (MSE), Structural-Similarity-Index-Measure (SSIM), and Peak Signal-to-Noise Ratio (PSNR) for the comparison of the ground-truth 4REPS to a single repetition (Original), images generated from DnCNN (Original: DnCNN) and 7TCDM (Original:7TCDM) (*: p < 0.05, **: p < 0.01, ***: p < 0.001) over the ADRC cohort of 19 subjects, 6 2D slices each, with each of the 114 slices depicted as a separate point.

### 3.2 Rating Assessment

Raters provided blinded assessments (1 being the highest score, and 5 the lowest) of all original, averaged, and denoised images (**Fig. 4**), across 6 categories of image quality interpretation.

**Figure 4.**
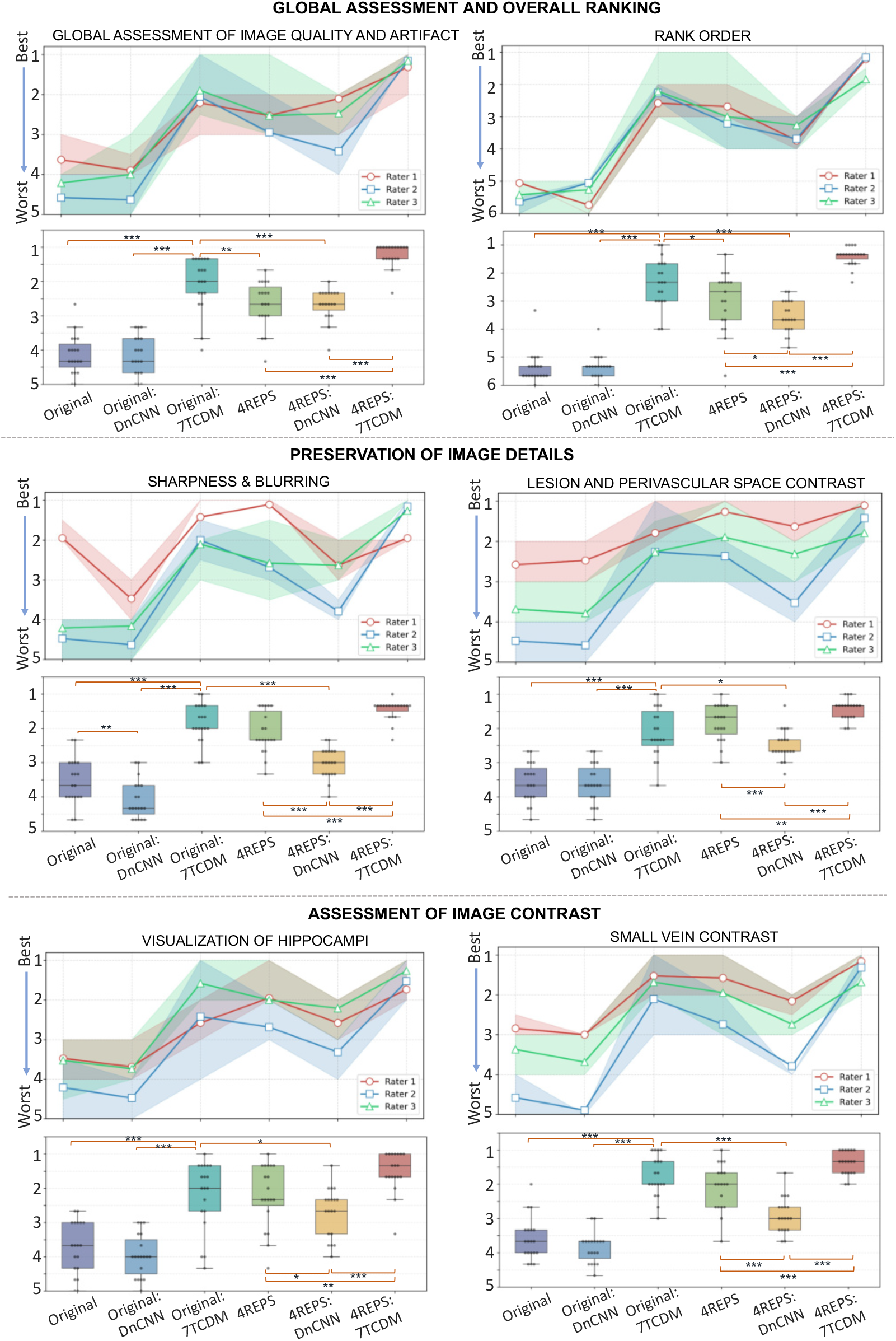
Box plots of the three neuroradiologists’ ratings (first row) on all 19 test subjects across 6 different image contrasts (Original, Original: DnCNN, Original:7TCDM, 4REPS, 4REPS: DnCNN and 4REPS: 7TCDM) assessing three aspects of image quality (Global Assessment and Overall Ranking, Preservation of Image Details, and Assessment of Image Contrast). The second row displays the averaged ratings across all radiologists for each assessment component (*: p < 0.05, **: p < 0.01, ***: p < 0.001).

#### Global Assessment and Overall Ranking

4REPS:7TCDM was rated as the best global image quality (1.21±0.34), with Original:7TCDM as second highest (score: 2.05±0.78), significantly better than Original (4.14±0.58, *p<0.0001*) and Original:DnCNN images (4.18±0.57, *p<0.0001*), as well as 4REPS (2.67±0.68, *p=0.0032*) and 4REPS:DnCNN, 2.67±0.46, *p=0.0008*).

All raters highly agreed with the ranking of each method (Averaged Gwet’s AC^28^: 0.720). 4REPS:7TCDM ranked highest (p < 0.0001 for all) and Original:7TCDM was ranked second highest, higher than 4REPS (*p=0.0466*) and 4REPS:DnCNN (*p<0.0001*).

#### Preservation of Image Details (Sharpness and Visualization of hippocampi)

The Original:7TCDM model was rated as better than Original (*p<0.0001*) and Original:DnCNN (*p<0.0001*) but not significantly different to 4REPS (*p=0.0557*) on sharpness. Original:DnCNN (4.09±0.55) and 4REPS:DnCNN (3.02±0.44) were each rated as having more blurring than Original (3.54±0.67, *p=0.0062*) and 4REPS (2.12 ±0.60, *p<0.0001*), respectively.

For hippocampal visualization, Original:7TCDM (2.19 ±1.01) received significantly higher ratings than Original (3.74±0.92, *p<0.0001*) and Original:DnCNN (3.96±0.72, *p<0.0001*). Similarly, 4REPS:7TCDM (1.51±0.59) was rated higher than 4REPS (2.12±0.92, *p=0.0045*) and 4REPS:DnCNN (2.70±0.72, *p<0.0001*).

#### Assessment of Image Contrast

Original:7TCDM was rated as having higher contrast of T2 bright areas such as white matter lesions and perivascular spaces over Original (*p < 0.0001*) and Original:DnCNN (*p < 0.0001*). These T2 bright areas were also more visible in Original:7TCDM and 4REPS:7TCDM compared to other images (**Fig. 5A**). Original:7TCDM also exhibited significantly better small vein contrast than 4REPS:DnCNN (*p < 0.0001*), and it was statistically equivalent to 4REPS (*p = 0.0905*).

**Figure 5.**
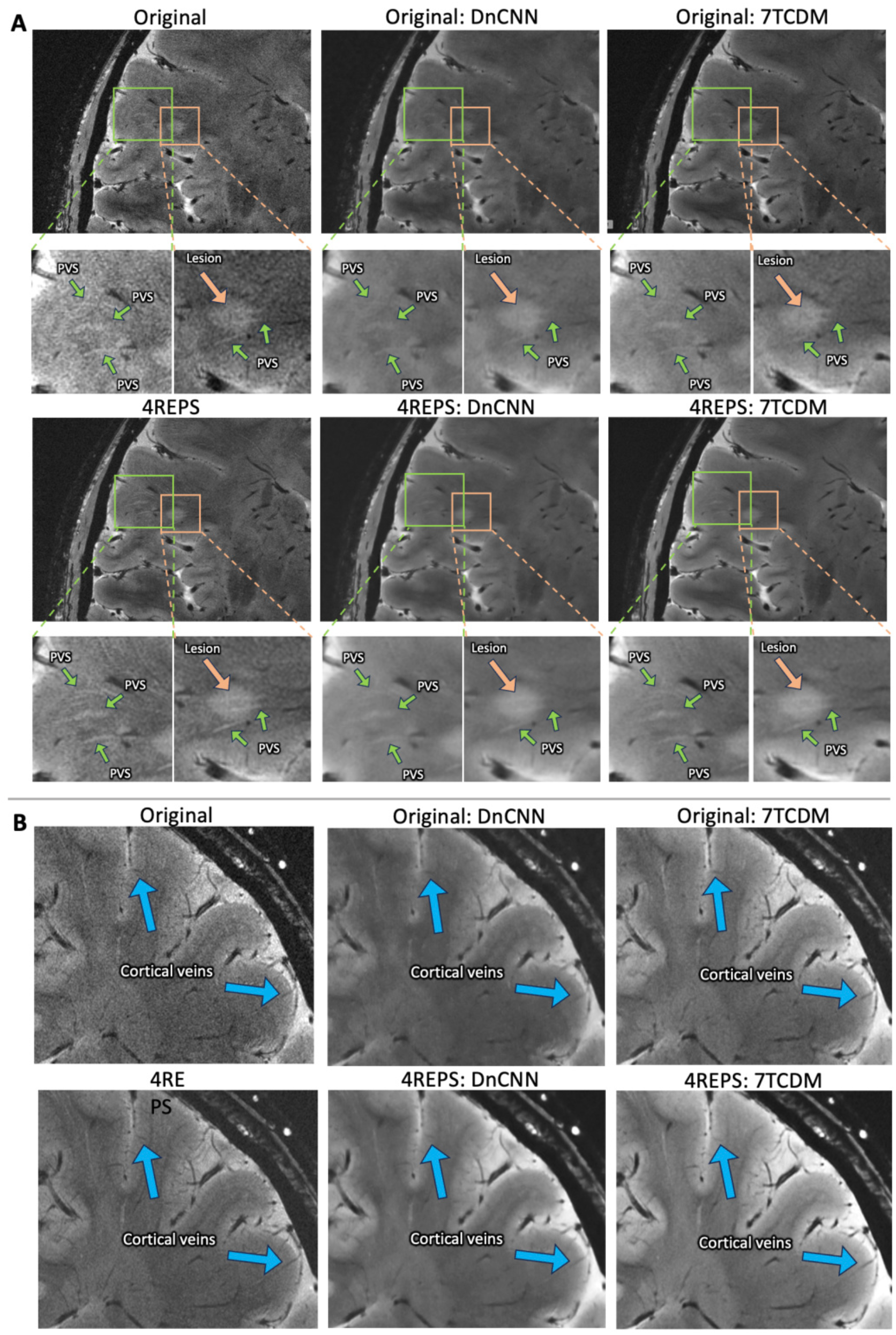
Visual comparison of image contrast preservation across original images and different methods for one example participant. **A.** Zoomed-in views of perivascular spaces (PVS, green arrows) and white matter hyperintensities (orange arrows). **B.** Small cortical veins (blue arrows).

Qualitative assessment of small cortical veins (**Fig. 5B**) showed sharp veins in Original:7TCDM but smoothed margins with Original:DnCNN and 4REPS:DnCNN, the latter two of which were statistically worse compared to 4REPS.

### 3.3 Generalization on 3D Multi-echo Gradient Echo MRI

We tested 7TCDM and DnCNN on 3D iron-sensitive multi-echo GRE (five echoes) to assess its generalizability to 7T GRE with different resolutions, contrast, and quality. **Figure 6A** shows one original image of three different echoes from an AD patient, alongside the denoised images generated by DnCNN and 7TCDM. Noise is less apparent in both reconstructions but at the cost of blurring for DnCNN. The R2* maps from the denoised DnCNN and 7TCDM models show reduced noise and more precise boundaries of deep gray nuclei in the basal ganglia (blue arrow, **Fig. 6A**) and midbrain (orange arrow, **Fig. 6A**). Both models were further tested on a public 7T multi-echo gradient echo dataset ^22^ (Siemens Healthineers, Germany), which covers primarily the parietal and occipital lobes (0.35 mm isotropic resolution, TR = 30 ms, and six echo times at 3.8, 7.6, 11.4, 15.2, 19.0, and 22.8 ms), subjectively improving image quality without any retraining (**Fig. 6B**).

**Figure 6.**
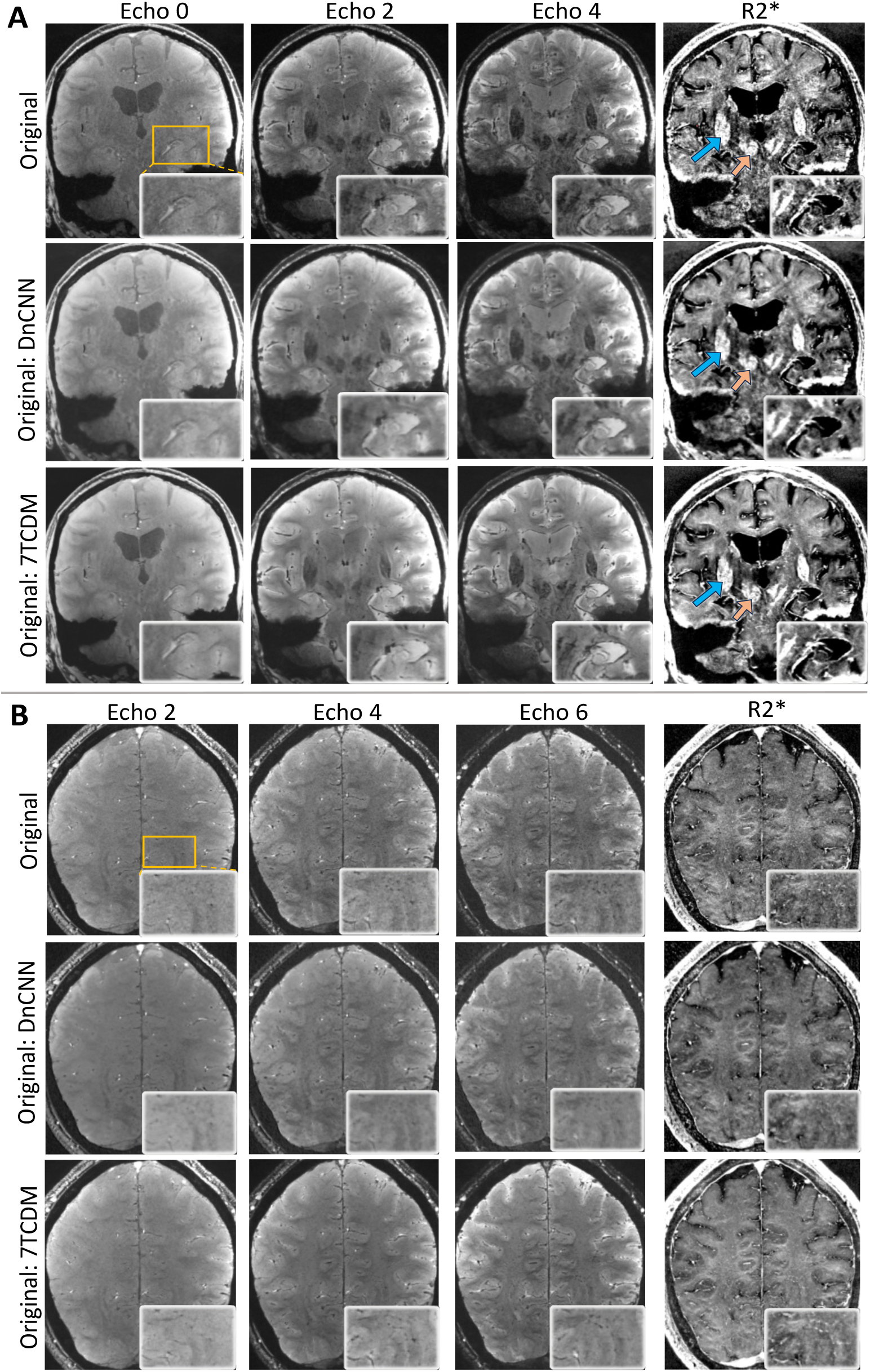
Models’ generalizability on 3D Multi-echo Gradient Echo MRIs from **(A)** our ADRC patients and **(B)** a public 7T dataset ^22^. DnCNN and 7TCDM models were tested on each echo without models retraining. The figure depicts matched slices from echo 0, 2, and 4 from one AD patient and echo 2, 4, and 6 from one participant from public data, alongside computed R2*. Top inset zooms in on the left hippocampus. Arrows pointing to putamen (blue) and red nuclei (orange).

## 4. Discussion

We present 7T Conditional Diffusion Model (7TCDM), a generative AI model for denoising 7T MRI. Individual short acquisitions served as a condition to guide the reverse diffusion process, enabling it to progressively remove noise and achieve a quality comparable to averaging four sequential short acquisitions. SSIM was integrated into the optimization process to enhance image contrast, aligning with clinical requirements for the diagnosis of finer structural changes and lesions. 7TCDM enables high-quality 7T MRI denoising from a single repetition, confirmed by both quantitative metrics and expert rater evaluation. It also generalizes to 3D GRE data with minimal preprocessing and no additional training.

Denoising methods using diffusion models have primarily been applied to 1.5T or 3T MRI. Some studies focus on super-resolution tasks, generating 7T-like images from 3T scans, particularly in diffusion MRI ^29^ or MR spectroscopic imaging ^30^. Others have explored recovering bone microstructure from CT images ^31^, removing noise from optical coherence tomography ^32^ and positron emission tomography ^19^, demonstrating the versatility of diffusion models across various imaging modalities. Commonly used methods for denoising 7T MRIs remain classical post-processing techniques, such as averaging multiple repetitions for 7T structural MRI ^23^, local complex Principal Component Analysis (PCA) ^14^ for quantitative MRI, and NOise Reduction with DIstribution Corrected (NORDIC) PCA for functional MRI ^15^. 7TCDM utilizes advanced generative AI techniques to achieve 7T structural MRI denoising without compromising key image features and anatomic details, which could pave the way for more efficient clinical MRI protocols.

Our study, like others ^31,33^, found that the commonly-used MSE/PSNR/SSIM are inadequate for clinical characterization, and rater assessments are revealing. While DnCNN achieved significantly higher MSE/PSNR/SSIM values compared to original images, it was evaluated lower by raters in all rating assessment metrics, likely due to smoothing of image details ^11^. In contrast, 7TCDM achieved improved MSE/PSNR/SSIM and rating assessments. Furthermore, 7TCDM denoising was generalized to both concurrently acquired and publicly available 3D multi-echo GRE, which were of different resolution and contrast, without any additional training. This can facilitate high-quality R2* mapping, which should propel qualitative and quantitative analysis of iron deposition in the brain ^3,34^ in Alzheimer’s disease and other neurodegenerative conditions. This generalization ability was supported by the model’s conditioning on original MRI data, making it applicable to various types of artifacts and noises. Specifically, while Gaussian noise was introduced and removed during the diffusion process, 7TCDM was conditioned on real images that contain Rician noise (e.g., in GRE), allowing it to learn corrections that reflect true acquisition-specific distortions.

Limitations include the small sample size of the training set, which might constrain the model’s capacity for denoising. However, we maximized data augmentation on the 2D slices to achieve robust results on the 19 ADRC held-out subjects even with the small dataset. The 7TCDM 2D model denoises slices independently, which may limit spatial consistency across slices. While it effectively preserves anatomical continuity and reduces some through-plane artifacts (**Supplementary Fig. 5**), full 3D models that leverage volumetric context could further improve spatial consistency. However, this would require more computational resources and larger training datasets.

Our future 7T work will include 7TCDM applied to various acquisitions, scanning conditions, and imaging contrasts, with contrast-specific training as necessary. Finally, we intend to compare 7TCDM with upcoming generative modeling paradigms, such as flow matching ^35^.

In conclusion, our Conditional Diffusion Model (7TCDM) enhances images from a single 5-minute gradient-echo (MGE) acquisition, superior to convolutional neural networks. These advancements could lead to more efficient MRI protocols not only at 7T but also at 3T and 1.5T.

## Supplementary material

**Supplementary Table 1.** Rater scoring scale for image quality assessment.

|  | GLOBAL<br>ASSESSMENT OF<br>IMAGE QUALITY<br>AND ARTIFACT | SHARPNESS<br>/ BLURRING | VISUALIZATION<br>OF HIPPOCAMPI | LESION AND<br>PERIVASCULAR<br>SPACE<br>CONTRAST | SMALL VEIN<br>CONTRAST |
| --- | --- | --- | --- | --- | --- |
| Description | Overall clarity, visibility of anatomical structures, and general impression. | Sharpness and clarity of brain structures, without blurring or loss of detail. | The sharpness and detail of the hippocampal structures, ensuring they are clearly identifiable without blurring or loss of detail | Contrast and clarity of the lesion and perivascular space | Contrast and clarity of small vein |
| Score | 1=excellent - no noise and no artifact | 1=excellent - no loss of anatomical detail and no blurring | 1=excellent | 1=excellent - no loss of lesion and perivascular space | 1=excellent - no loss of small vein contrast |
|  | 2=very good - limited noise and artifact | 2=very good - minimal loss of detail and/or minimal blurring | 2=very good | 2=very good - minimal loss of lesion and perivascular space | 2=very good - minimal loss of small vein contrast |
|  | 3=good - mild noise and artifact | 3=good - mild loss of anatomical detail and/or mild blurring | 3=good | 3=good - mild loss of lesion and perivascular space | 3=good - mild loss of small vein contrast |
|  | 4=adequate - moderate noise and artifact | 4=adequate - moderate loss of detail and/or moderate blurring | 4=adequate | 4=adequate - moderate loss of lesion and perivascular space | 4=adequate - moderate loss of small vein contrast |
|  | 5= limited - noisy and artifactual | 5= limited - significant loss of anatomical detail and/or significant blurring | 5=limited | 5=limited - significant loss of lesion and perivascular space | 5=limited - significant loss of small vein contrast |

### Experimental Settings

7TCDM was trained on 11 scans from MRIs of five healthy adults for 50,000 iterations with *T*=10 (see Supplementary Ablation study) and a batch size of 8. We used an AdamW optimizer with betas set to (0.9, 0.999), a weight decay of 0.01 and a learning rate of 1×10^−4^. λ_SSIM_ was chosen as 0.5 (see Supplementary Ablation study). All experiments were conducted on a computing node with 4 NVIDIA A6000 GPUs, each with 48GB of memory. We used a Denoising Diffusion Implicit Model (DDIMs) sampler ^35^ for efficient sampling, where only fewer steps (<= *T*) were required to obtain the final denoised images (∼8s per 2D slice).

DnCNN was trained for 30 epochs with a batch size of 4, a learning rate of 1×10^−4^ and an Adam optimizer. The network depth was set as the default value of 17. For a fair comparison, we adopted the same data preprocessing methods as in 7TCDM including the histogram matching and data augmentation described in Section 2.2. The same SSIM was integrated into the default MSE loss function during training.

### Ablation Study

Inspired by Fast-DDPM ^45^, which demonstrates that a smaller training step *T* leads to better performance in denoising tasks, we examined the effect of *T* on model performance during training and inference in **Supplementary Fig.1** (first row). Reducing *T* to 10 significantly decreased inference time while simultaneously improving image quality, as measured by MSE, SSIM, and PSNR. This reduction in time steps led to a more efficient model with faster inference times without compromising the quality of the reconstructed images.

We evaluated the impact of weight λ_ssim_on the performance of both 7TCDM and DnCNN (**Supplementary Fig.1**, second row). A large SSIM improvement accompanied increasing λ_ssim_ by 7TCDM, which was at the cost of a small increase in MSE and decrease PSNR. All three metrics were decreased on DnCNN when λ_ssim_ was increased from 0 to 0.5. The model with λ_ssim_ = 0.5 was selected as the best choice, prioritizing SSIM improvement and lesion contrast preservation (**Supplementary Fig.2**) with minimal sacrifice in other metrics for 7TCDM. Experiments showed λ_ssim_ = 1 led to unstable optimization and unsuccessfully training so it was not included in our ablation study.

**Supplementary Figure 1.**
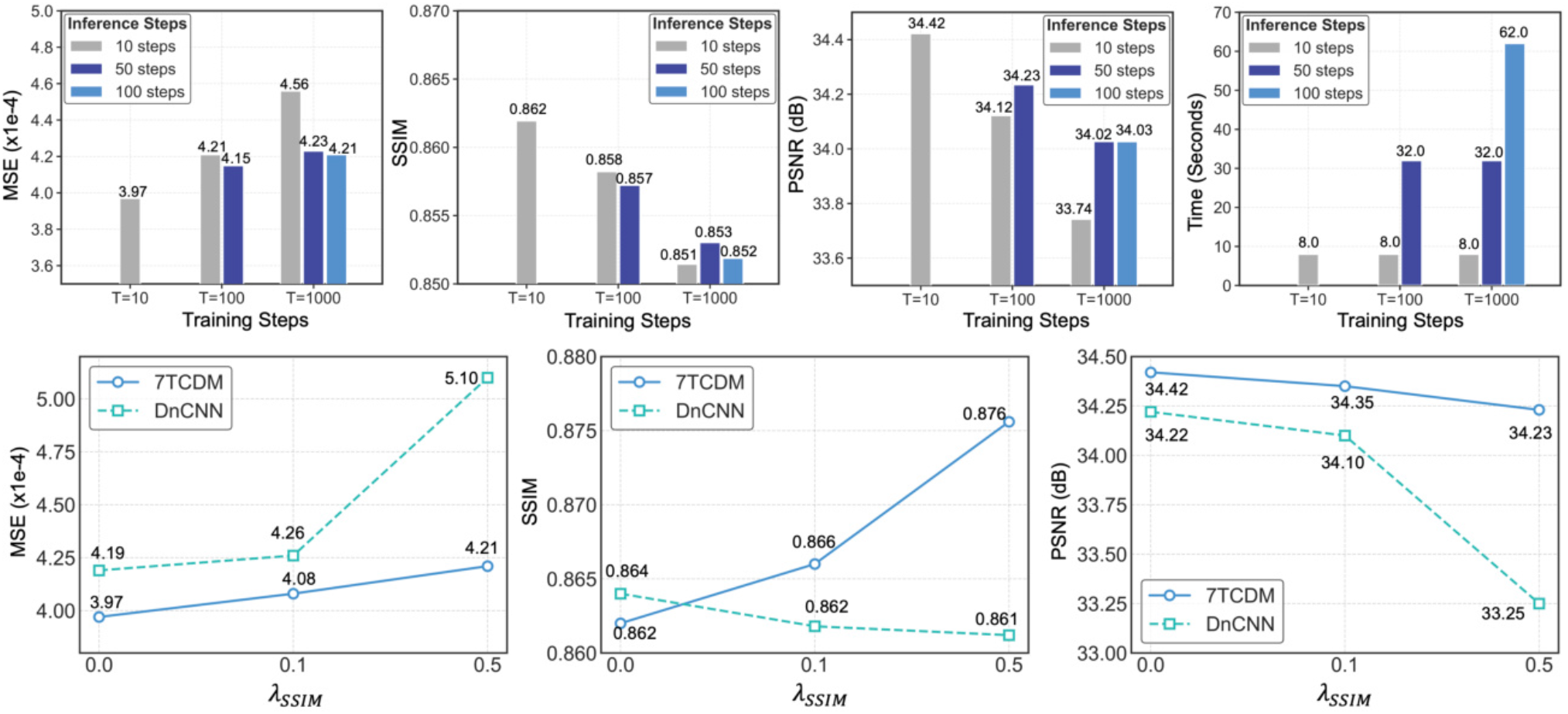
Ablation study on the impact of training steps, inference steps, and the weight of the SSIM term (λ_ssim_) on the model’s performance. The top row shows the effect of different inference steps (10, 50, 100 steps) and training steps (T = 10, 100, 1000) on MSE, SSIM, PSNR, and inference time. The bottom row compares the performance of both 7TCDM and DnCNN models with varying impacts of SSIM weighting (λ_ssim_= 0, 0.1, 0.5).

**Supplementary Figure 2.**
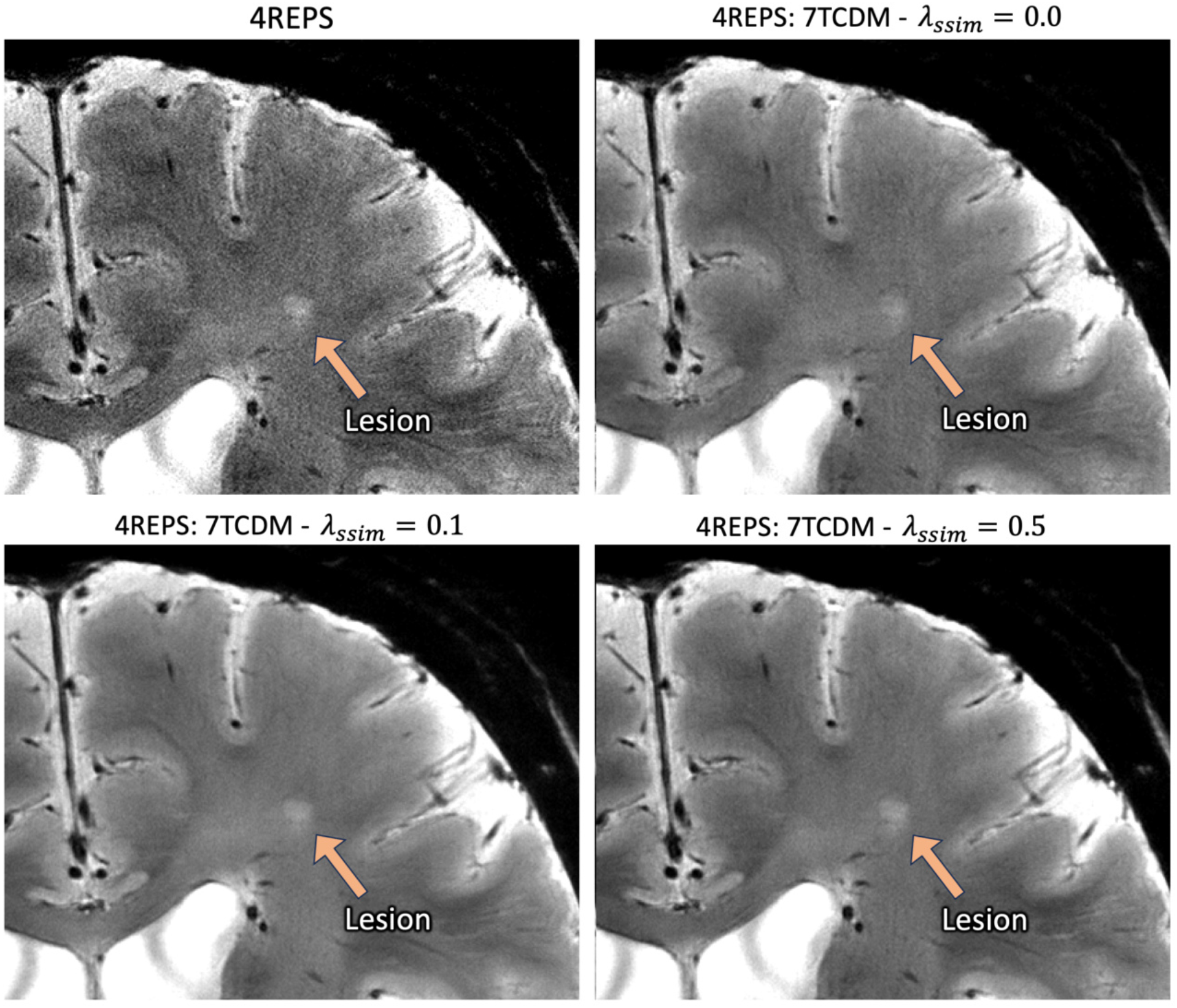
Comparison of lesion visibility in images reconstructed with different λ_ssim_ values using the 7TCDM model.

### Residual Comparisons

The residuals between the original images and the averaged (4REPS) and denoised images visualize the information that is removed from the averaging/denoising process. DnCNN residuals subjectively show more anatomic detail, while the 7TCDM residual image contains relatively less structural detail (**Supplementary Fig.3**). We quantified this by computing the SSIM between these residual images and the ground truth 4REPS: the SSIM for 7TCDM residuals was 0.0905 ± 0.0109, whereas the SSIM for DnCNN residuals was higher at 0.1160 ± 0.0096. This reflects a higher similarity between 4REPS and DnCNN residuals, indicating that more anatomic information is present in DnCNN residuals and thus “removed” in the DnCNN reconstructions.

**Supplementary Figure 3.**
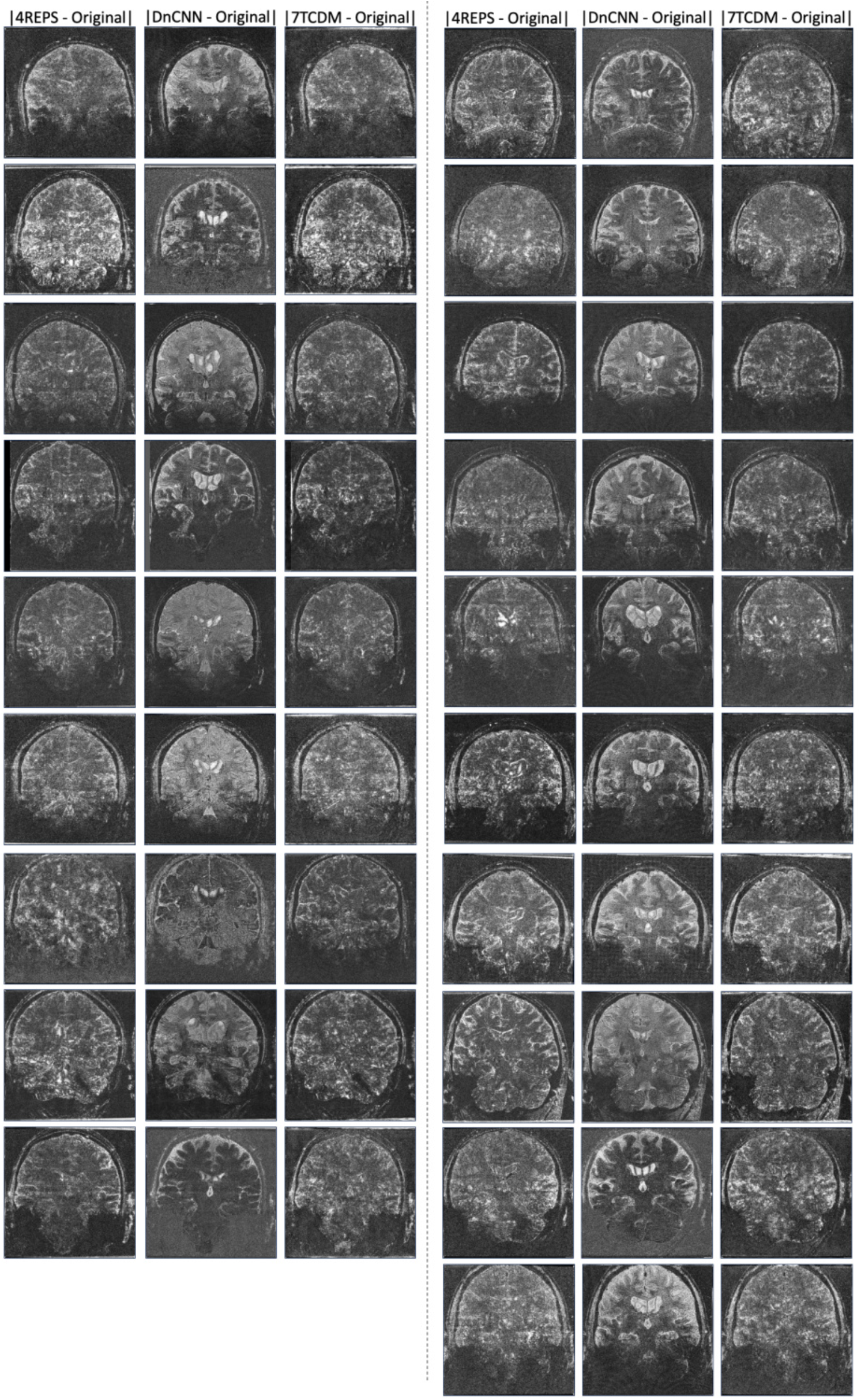
Visualization of difference (residual) maps between original image and averaged (4REPS) / denoised images (Original:DnCNN and Original:7TCDM) from all 19 participants. DnCNN difference maps show significant residual structural information, more so than 7TDCM and 4REPS. |·| means absolute value.

**Supplementary Figure 4.**
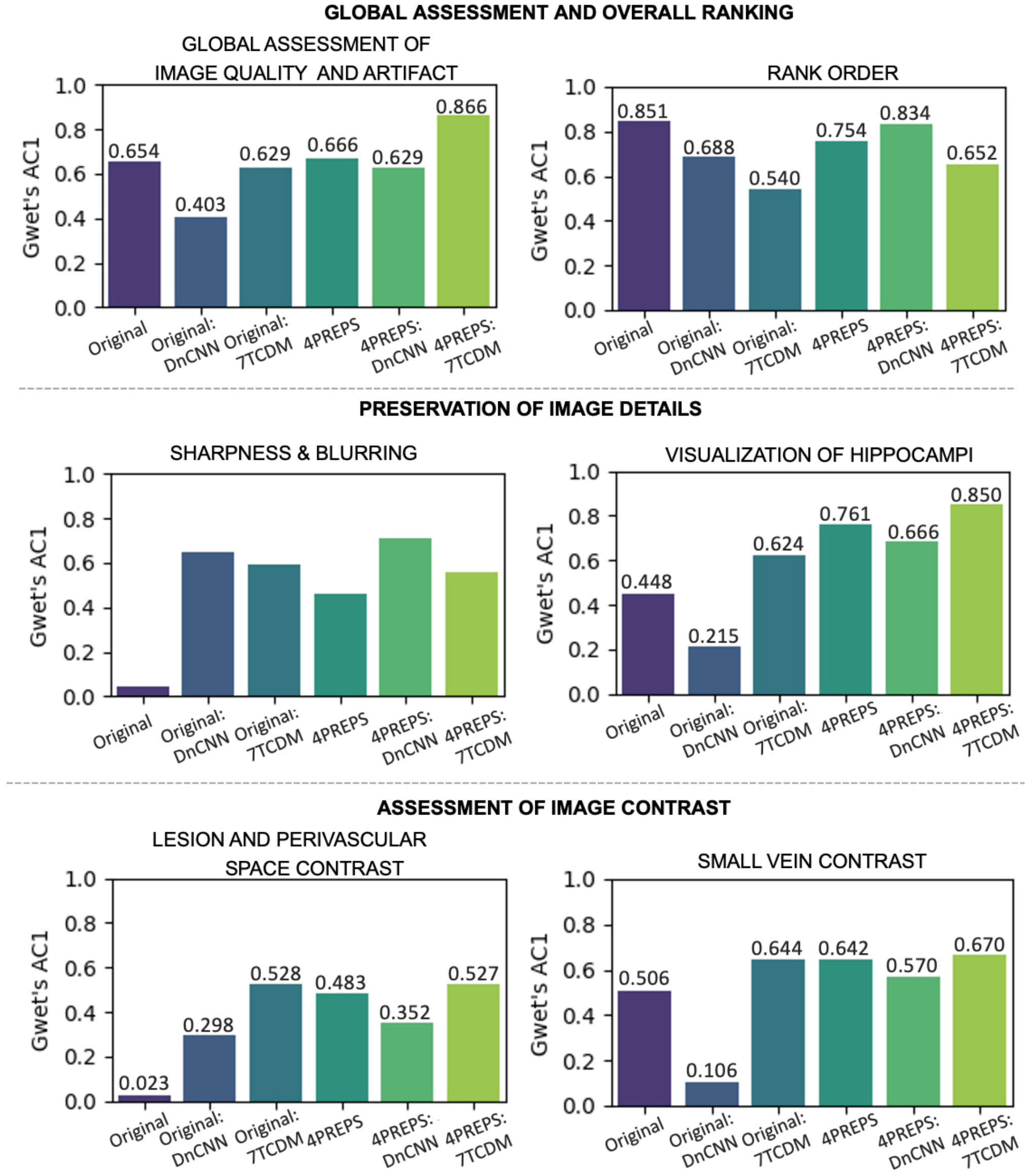
Rater agreement evaluated by Gwet’s AC1 scores for various image quality assessments. Higher AC1 scores indicate better agreement among three raters.

**Supplementary Figure 5.**
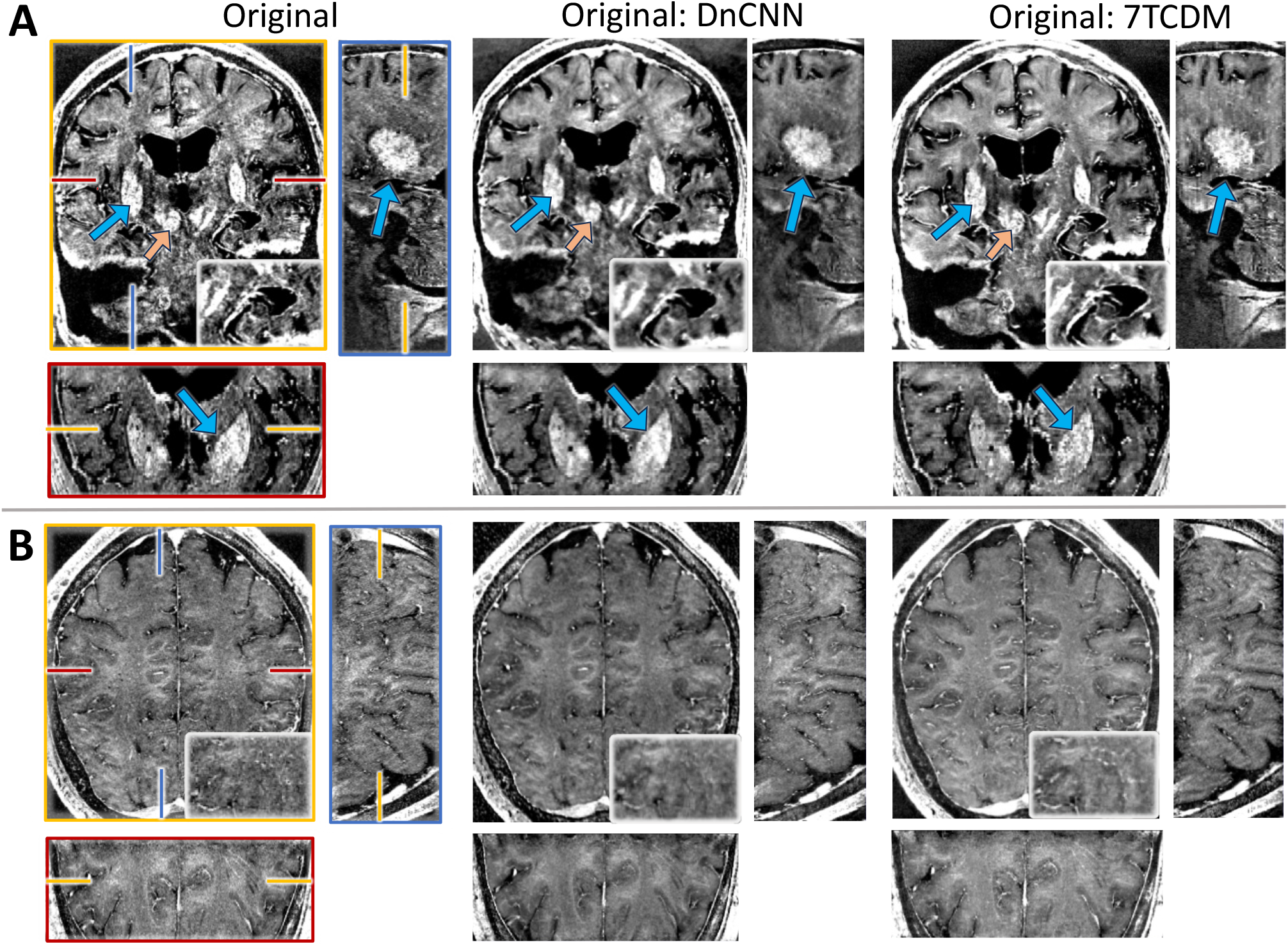
Computed R2* from **(A)** one AD patient and **(B)** one participant from public data in three planes. Arrows pointing to the putamen (blue) and red nuclei (orange).

## Reference

1. Okada T, Fujimoto K, Fushimi Y, et al. Neuroimaging at 7 Tesla: a pictorial narrative review. Quant Imaging Med Surg. 2022;12(6):3406–3435. doi:10.21037/qims-21-969

2. Perera Molligoda Arachchige AS, Garner AK. Seven Tesla MRI in Alzheimer’s disease research: State of the art and future directions: A narrative review. AIMS Neurosci. 2023;10(4):401–422. doi:10.3934/Neuroscience.2023030

3. Zeineh MM, Chen Y, Kitzler HH, Hammond R, Vogel H, Rutt BK. Activated iron-containing microglia in the human hippocampus identified by magnetic resonance imaging in Alzheimer disease. Neurobiol Aging. 2015;36(9):2483–2500. doi:10.1016/j.neurobiolaging.2015.05.022

4. Karamat M, Darvish-Molla S, Santos Diaz A. Opportunities and Challenges of 7 Tesla Magnetic Resonance Imaging: A Review. Crit Rev Biomed Eng. 2016;44. doi:10.1615/CritRevBiomedEng.2016016365

5. Gallichan D, Marques JP. Optimizing the acceleration and resolution of three-dimensional fat image navigators for high-resolution motion correction at 7T. Magn Reson Med. 2017;77(2):547–558. doi:10.1002/mrm.26127

6. Bazin PL, Nijsse HE, van der Zwaag W, et al. Sharpness in motion corrected quantitative imaging at 7T. NeuroImage. 2020;222:117227. doi:10.1016/j.neuroimage.2020.117227

7. DiGiacomo P, Maclaren J, Aksoy M, et al. A within-coil optical prospective motion-correction system for brain imaging at 7T. Magn Reson Med. 2020;84(3):1661–1671. doi:10.1002/mrm.28211

8. Zhang K, Zuo W, Chen Y, Meng D, Zhang L. Beyond a Gaussian Denoiser: Residual Learning of Deep CNN for Image Denoising. Trans Img Proc. 2017;26(7):3142–3155. doi:10.1109/TIP.2017.2662206

9. Cheng H, Vinci-Booher S, Wang J, et al. Denoising diffusion weighted imaging data using convolutional neural networks. PloS One. 2022;17(9):e0274396. doi:10.1371/journal.pone.0274396

10. Zormpas-Petridis K, Tunariu N, Curcean A, et al. Accelerating Whole-Body Diffusion-weighted MRI with Deep Learning–based Denoising Image Filters. Radiol Artif Intell. 2021;3(5):e200279. doi:10.1148/ryai.2021200279

11. Kazerouni A, Aghdam EK, Heidari M, et al. Diffusion models in medical imaging: A comprehensive survey. Med Image Anal. 2023;88:102846. doi:10.1016/j.media.2023.102846

12. Bagley BA, Petrov S, Cheng G, et al. Generative Editing via Convolutional Obscuring (GECO): A Generative Adversarial Network for MRI de-artifacting. Published online February 14, 2023:2022.09.21.22280206. doi:10.1101/2022.09.21.22280206

13. Arjovsky M, Bottou L. Towards Principled Methods for Training Generative Adversarial Networks. In:; 2017. Accessed April 23, 2025. https://openreview.net/forum?id=Hk4_qw5xe

14. Bazin PL, Alkemade A, van der Zwaag W, Caan M, Mulder M, Forstmann BU. Denoising High-Field Multi-Dimensional MRI With Local Complex PCA. Front Neurosci. 2019;13:1066. doi:10.3389/fnins.2019.01066

15. Vizioli L, Moeller S, Dowdle L, et al. Lowering the thermal noise barrier in functional brain mapping with magnetic resonance imaging. Nat Commun. 2021;12(1):5181. doi:10.1038/s41467-021-25431-8

16. Tsuji N, Kobayashi T, Ueda J, Saito S. Development and Evaluation of Deep Learning-Based Reconstruction Using Preclinical 7T Magnetic Resonance Imaging. Appl Sci. 2023;13(11):6567. doi:10.3390/app13116567

17. Ho J, Jain A, Abbeel P. Denoising diffusion probabilistic models. In: Proceedings of the 34th International Conference on Neural Information Processing Systems. NIPS ‘20. Curran Associates Inc.; 2020:6840–6851.

18. Ma J, Zhu Y, You C, Wang B. Pre-trained Diffusion Models for Plug-and-Play Medical Image Enhancement. In: Greenspan H, Madabhushi A, Mousavi P, et al., eds. Medical Image Computing and Computer Assisted Intervention – MICCAI 2023. Springer Nature Switzerland; 2023:3–13. doi:10.1007/978-3-031-43898-1_1

19. Gong K, Johnson K, El Fakhri G, Li Q, Pan T. PET image denoising based on denoising diffusion probabilistic model. Eur J Nucl Med Mol Imaging. 2024;51(2):358–368. doi:10.1007/s00259-023-06417-8

20. Wang J, Levman J, Pinaya WHL, Tudosiu PD, Cardoso MJ, Marinescu R. InverseSR: 3D Brain MRI Super-Resolution Using a Latent Diffusion Model. Published online August 23, 2023. Accessed October 6, 2024. http://arxiv.org/abs/2308.12465

21. Xiang T, Yurt M, Syed A, Setsompop K, Chaudhari A. DDM$^2$: Self-Supervised Diffusion MRI Denoising with Generative Diffusion Models.; 2023. doi:10.48550/arXiv.2302.03018

22. Dresbach S, Huber R, Gülban ÖF, et al. Characterisation of laminar and vascular spatiotemporal dynamics of CBV and BOLD signals using VASO and ME-GRE at 7T in humans. Imaging Neurosci. 2024;2:1–16. doi:10.1162/imag_a_00263

23. DiGiacomo P, Maclaren J, Aksoy M, et al. A within-coil optical prospective motion-correction system for brain imaging at 7T. Magn Reson Med. 2020;84(3):1661–1671. doi:10.1002/mrm.28211

24. Jenkinson M, Bannister P, Brady M, Smith S. Improved Optimization for the Robust and Accurate Linear Registration and Motion Correction of Brain Images. NeuroImage. 2002;17(2):825–841. doi:10.1006/nimg.2002.1132

25. Nichol AQ, Dhariwal P. Improved Denoising Diffusion Probabilistic Models. In: Proceedings of the 38th International Conference on Machine Learning. PMLR; 2021:8162–8171. Accessed November 1, 2024. https://proceedings.mlr.press/v139/nichol21a.html

26. Wang Z, Bovik AC, Sheikh HR, Simoncelli EP. Image quality assessment: from error visibility to structural similarity. IEEE Trans Image Process. 2004;13(4):600–612. doi:10.1109/TIP.2003.819861

27. Song J, Meng C, Ermon S. DENOISING DIFFUSION IMPLICIT MODELS. Published online 2021.

28. Gwet KL. Computing inter-rater reliability and its variance in the presence of high agreement. Br J Math Stat Psychol. 2008;61(Pt 1):29–48. doi:10.1348/000711006X126600

29. Zhu X, Zhang W, Li Y, O’Donnell LJ, Zhang F. When Diffusion MRI Meets Diffusion Model: A Novel Deep Generative Model for Diffusion MRI Generation. In: Linguraru MG, Dou Q, Feragen A, et al., eds. Medical Image Computing and Computer Assisted Intervention – MICCAI 2024. Springer Nature Switzerland; 2024:530–540. doi:10.1007/978-3-031-72069-7_50

30. Dong S, Cai Z, Hangel G, et al. A Flow-based Truncated Denoising Diffusion Model for super-resolution Magnetic Resonance Spectroscopic Imaging. Med Image Anal. 2025;99:103358. doi:10.1016/j.media.2024.103358

31. Chan TJ, Rajapakse CS. A Super-Resolution Diffusion Model for Recovering Bone Microstructure from CT Images. Radiol Artif Intell. 2023;5(6):e220251. doi:10.1148/ryai.220251

32. Hu D, Tao YK, Oguz I. Unsupervised denoising of retinal OCT with diffusion probabilistic model. In: Medical Imaging 2022: Image Processing. Vol 12032. SPIE; 2022:25–34. doi:10.1117/12.2612235

33. Sampat MP, Wang Z, Gupta S, Bovik AC, Markey MK. Complex Wavelet Structural Similarity: A New Image Similarity Index. IEEE Trans Image Process. 2009;18(11):2385–2401. doi:10.1109/TIP.2009.2025923

34. Tran D, DiGiacomo P, Born DE, Georgiadis M, Zeineh M. Iron and Alzheimer’s Disease: From Pathology to Imaging. Front Hum Neurosci. 2022;16:838692. doi:10.3389/fnhum.2022.838692

35. Lipman Y, Chen RTQ, Ben-Hamu H, Nickel M, Le M. Flow Matching for Generative Modeling. In:; 2022. Accessed April 3, 2025. https://openreview.net/forum?id=PqvMRDCJT9t

